# Assessment of NR4A Ligands that Directly Bind and Modulate the Orphan Nuclear Receptor Nurr1

**DOI:** 10.1101/2020.05.22.109017

**Authors:** Paola Munoz-Tello, Hua Lin, Pasha Khan, Ian Mitchelle S. de Vera, Theodore M. Kamenecka, Douglas J. Kojetin

## Abstract

Nurr1/NR4A2 is an orphan nuclear receptor transcription factor implicated as a potential drug target for neurological disorders including Alzheimer’s and Parkinson’s diseases. Previous studies identified small molecule modulators of NR4A nuclear receptors including Nurr1 and Nur77/NR4A1; it remains unclear whether these ligands affect Nurr1 through direct binding or indirect non-binding mechanisms. We assessed a panel of twelve ligands reported to affect NR4A activity for Nurr1-dependent and Nurr1-independent transcriptional effects and binding to the Nurr1 ligand-binding domain (LBD). Most of the NR4A ligands show Nurr1-independent effects on transcription in a cell type-specific manner, suggesting they may function through binding to effector proteins whose downstream activities influence Nurr1 function. Protein NMR spectroscopy structural footprinting data show that 4-amino-7-chloroquinoline derivatives (amodiaquine and chloroquine) and cytosporone B directly bind the Nurr1 LBD. In contrast, other NR4A ligands including commercially available compounds such as C-DIM12, celastrol, camptothecin, IP7e, isoalantolactone, and TMPA do not bind the Nurr1 LBD. Interestingly, previous crystal structures indicate that cytosporone B analogs bind to surface pockets in the Nur77 LBD, but protein NMR data indicate cytosporone B likely binds to the Nurr1 orthosteric pocket. These findings should influence medicinal chemistry efforts that desire to optimize Nurr1-binding ligands as opposed to ligands that function through binding to Nurr1 effector proteins.

## INTRODUCTION

The NR4A nuclear receptor transcription factors regulate important physiological processes including cellular homeostasis, metabolic regulation, apoptosis, and cell differentiation (*1*). The three members of the NR4A family—NR4A1 (Nur77), NR4A2 (Nurr1), and NR4A3 (NOR1)—regulate the transcription of target genes through binding to specific DNA response element sequences, including a monomeric NGFI-B Response Element (NBRE) motif and a Nur-RXR heterodimer Response Element (NurRE) motif. Nuclear receptors are generally classified as ligand-dependent transcription factors. However, the NR4As have an unconventional orthosteric ligand-binding pocket compared to other nuclear receptors. Crystal structures of the Nur77 and Nurr1 ligand-binding domains (LBDs) show a collapsed orthosteric pocket that is filled with residues containing bulky hydrophobic sidechains suggesting these receptors may function independent of binding ligand within an orthosteric pocket (*2, 3*). Furthermore, contributing to their status as orphan receptors, it remains unclear if the NR4As are regulated by binding physiological or endogenous ligands, although unsaturated fatty acids (*4-6*) and dopamine metabolites (*7*) have been shown to interaction with the Nur77 and/or Nurr1 LBDs.

Regulating Nurr1 activity with small molecule activating ligands (agonists) is implicated to provide a therapeutic benefit in several Nurr1-related diseases including neurological disorders, inflammation, autoimmunity, cancer, and multiple sclerosis (*8-10*). To develop therapies to treat these diseases, several groups have initiated studies to discover small molecules that modulate Nurr1 transcription. Among these studies, compounds with a 4-amino-7-chloroquinoline scaffold—including amodiaquine, chloroquine, and glafenine—were identified in a high-throughput screen as ligands that increase Nurr1 transcription in human neuroblastoma SK-*N*-BE(2)-C cells (*11*). Amodiaquine improves behavioral alterations in a Parkinson’s disease animal model (*11*) and improves neuropathology and memory impairment in an Alzheimer’s disease animal model (*12*).

Amodiaquine is the most potent and efficacious Nurr1 agonist of the 4-amino-7-chloroquinoline compounds, but these compounds are not Nurr1 specific; they also target other proteins including apelin receptor (*13*) and are capable of antiviral (*14, 15*) and antimalarial (*16*) activity. However, knowledge that the 4-amino-7-chloroquinoline scaffold can directly bind to the Nurr1 LBD opens the path to future structure-activity relationship (SAR) studies to develop more potent and efficacious ligands with better specificity for Nurr1 over other molecular targets. Screening efforts have identified other classes of Nurr1-activating ligands with poorly defined mechanisms of action, and ligands reported to influence Nur77 activity represent another potential source to discover Nurr1 ligands since evolutionarily related nuclear receptors often display broad specificity for similar ligands. This is true for endogenous ligands—e.g., phospholipids for LRH-1 and SF-1, estrogen for ERα and ERβ—as well as synthetic ligands.

It would be useful to know if the Nurr1- and Nur77-activating compounds reported in the literature affect Nurr1 activity through direct binding to Nurr1, or via “off-target” mechanisms through binding to upstream effector proteins whose downstream activities influence Nurr1 function. We tested twelve ligands reported to affect the activity of Nurr1 or Nur77 for Nurr1-dependent transcription using cellular reporter assays and for direct binding to the Nurr1 LBD using a protein NMR structural foot-printing assay that provides information on ligand binding epitopes. We found that most of the ligands display cell type-specific transcriptional activities through Nurr1-dependent and Nurr1-independent mechanisms. Furthermore, using protein NMR we found that only three of the ligands directly bind to the Nurr1 LBD: amodiaquine, chloroquine, and cytosporone B. These findings should be of interest to medicinal chemists that want to focus on the discovery and optimization of Nurr1-binding ligands as opposed to ligands that affect Nurr1 activity through other indirect mechanisms.

## RESULTS

### NR4A ligand selection and properties

The twelve ligands we selected represent most, if not all, of the chemical scaffolds reported to modulate Nurr1- or Nur77-dependent transcription or other activities (**Figure 1**). Of these, nine were available from commercial sources and three others (SR10098, SR24237, and SR10658; compounds **1, 2**, and **3**, respectively) were synthesized in-house.

**Fig. 1.**
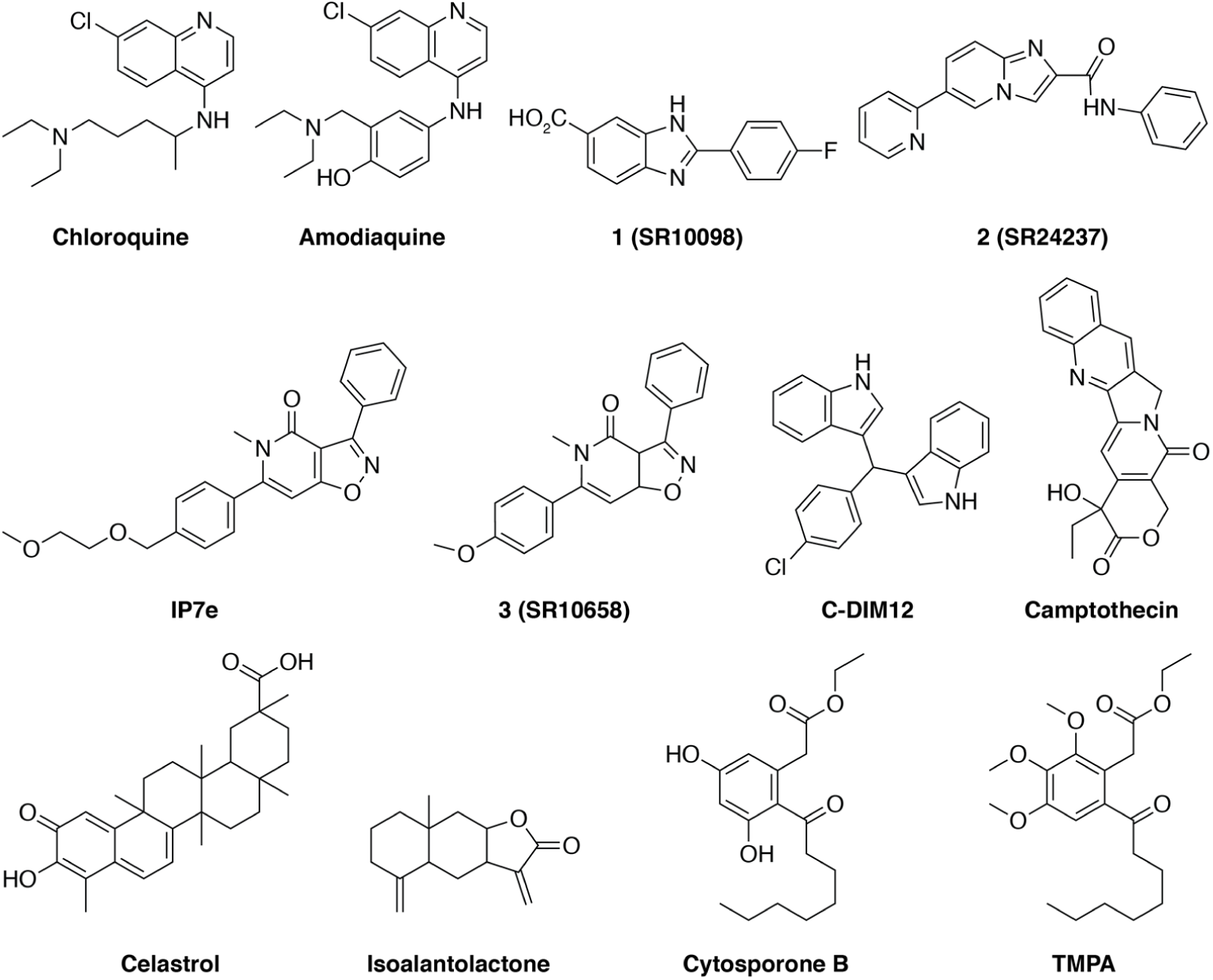
Chemical structures of the twelve NR4A ligands characterized in this study.

Amodiaquine and chloroquine are antimalarial ligands that contain a 4-amino-7-chloroquinoline scaffold and were identified in a screen using SK-*N*-BE(2)-C cells as activators of Nurr1 transcription with micromolar potency that directly bind to the Nurr1 LBD (*11*).

Four ligands were hits or optimized from a hit identified in different high-throughput screens as compounds that increase Nurr1-dependent transcription with low nanomolar potency using a luciferase reporter assay. The benzimidazole SR10098 (*17*) and the isoxazolo-pyridinone SR10658 (*18*) were discovered in a screen using the MN9D dopaminergic cellular model; IP7e is an analog of SR10658 reported to have improved solubility for *in vivo* studies and displayed *in vivo* efficacy in the experimental autoimmune encephalomyelitis mice (EAE) model of multiple sclerosis (MS) (*19*). The imidazopyridine SR24237 is an optimized compound from a hit discovered in two screens using Chinese hamster ovary (CHO) and a mouse neuronal N2A cell lines (*20*). An analog of SR24237 was reported to display neuroprotective and anti-inflammatory activity.

1,1-bis(3’-indolyl)-1-(*p*-chlorophenyl) methane, or C-DIM12, was reported as a synthetic Nurr1 activator that increases Nurr1 transcription, affects the expression of dopaminergic genes in pancreatic cells, keratinocyte epidermal cells, and primary neurons, and displays *in vivo* efficacy in models of Parkinson’s disease (*21-23*).

Camptothecin, an antitumor chemotherapeutic agent and cyclooxygenase-2 inhibitor, is a natural product identified in a high-throughput screen as a potent inhibitor of Nurr1 transcription that triggers an antitumor response by reducing Foxp3^+^ T regulatory cells and inducing IFNγ^+^ T helper 1 cells indicating Nurr1 may be a target for cancer immunotherapy (*24*).

Celastrol is a natural product that binds the Nur77 LBD with high affinity and influences Nur77 activity through multiple mechanisms (*25*). Nur77 and AMPKα are known to be involved in adipogenesis regulation (*26, 27*).

Isoalantolactone is a natural product discovered in a screen for ligands that inhibit Nur77 transcription and activate AMPKα in 3T3-L1 cells leading to a cascade of events that highlights a role for targeting Nur77 in protection against metabolic disorders and obesity (*28*).

Cytosporone B (CsnB) is a natural product identified as a Nur77 agonist that binds to the Nur7 LBD and enhances interaction of transcriptional coregulator proteins (*29*). CsnB was also shown to function as a Nurr1 agonist in BGC-823 human gastric carcinoma cells. Later work, using a chemical screen, identified an analog of CsnB called ethyl 2-[2,3,4-tri-methoxy-6-(1-octanoyl)phenyl]acetate, or TMPA, that showed low nanomolar affinity for the Nur77 LBD (*30*). Unlike CsnB, TMPA does not function as a canonical agonist of Nur77 transcription; rather, it inhibits the interaction between Nur77 and Liver kinase B1 (LKB1). Another CsnB analog that was not commercially available, PDNPA, inhibits the interaction between Nur77 and the MAP kinase p38α (*31*), suggesting inhibition of Nur77-kinase interactions may be a general mode of action for these compounds.

### Ligands display cell type-specific effects on Nurr1 transcription

To assess the effect of the compounds on Nurr1-mediated transcription, we transfected cells with a full-length Nurr1 expression plasmid along with one of two reporter plasmids either containing three copies of the NBRE or NurRE followed by the firefly luciferase gene. We performed the assays in three cell lines including HEK293T, a kidney embryonic cell line commonly used to assess general nuclear receptor activity, as well as two cell lines relevant to Nurr1 functions in neurons: PC12, a rat pheochromocytoma cell line exhibiting neuronal-like characteristics; and SK-*N*-BE(2)-C, a neuroblastoma cell line displaying moderate levels of tyrosine hydroxylase activity and dopamine-b-hydroxylase activity. Overall, we found that the ligands showed differential (cell type- or reporter-specific) to negligible activity in the various assays. We used four ligand concentrations in the assays that differ among all the ligands, which we based on previously reported cellular potencies in the original reports of the compounds as well as initial dose-response studies where we excluded higher concentrations that gave bell shaped response curves indicative of colloidal aggregation.

Amodiaquine (**Figure 2A**) and chloroquine (**Figure 2B**) increased activity of both luciferase reporters in SK-*N*-BE(2)-C cells consistent with previously published data (*11*); however, no activity and potentially a slight decrease in luciferase signal was observed for chloroquine in HEK293T and PC12 cells. SR10098 (**Figure 2C**) increased luciferase activity moderately in HEK293T cells, more efficaciously in SK-*N*-BE(2)-C cells, and showed no effect in PC12 cells. SR24237 (**Figure 2D**), and SR10658 (**Figure 2E**) showed dose-responsive increased activity in most conditions. IP7e (**Figure 2F**) only showed increased activity at the highest concentration tested (100 nM) in all conditions. C-DIM12 (**Figure 2G**) showed decreased activity in HEK293T and SK-*N*-BE(2)-C cells, but no effect in PC12 cells. Camptothecin (**Figure 2H**) showed decreased activity in all conditions except in PC12 cells with the NurRE reporter. Celastrol (**Figure 2I**) showed increased or decreased activity in a cell type-specific and DNA response element-specific manner. Isoalantolactone (**Figure 2J**) showed decreased activity in HEK293T and SK-*N*-BE(2)-C cells, but not activity in PC12 cells. Cytosporone B (**Figure 2K**) showed decreased activity for some conditions at higher concentrations, whereas TMPA (**Figure 2L**) showed no activity.

**Fig. 2.**
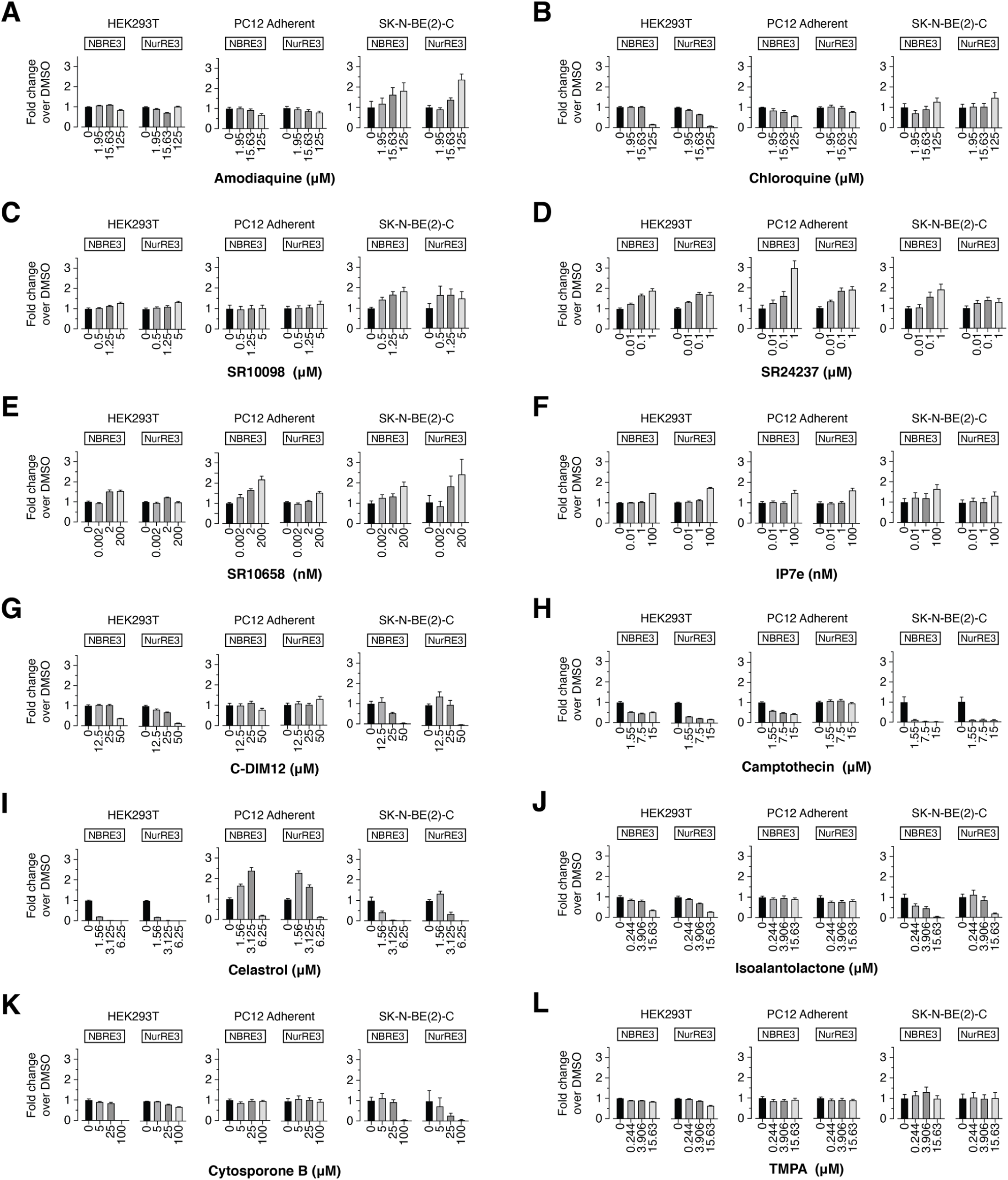
Effect of the twelve NR4A ligands on full-length Nurr1 transcription of a monomeric NBRE3 and RXR heterodimeric NurRE3 luciferase reporters in HEK293T, PC12, and SK-N-BE(2)-C cells (n=4, mean ± s.e.m; representative of ≥ 2 independent experiments).

### Ligand transcriptional effects independent of Nurr1

We performed two cellular assays to determine the specificity of the NR4A compounds in affecting Nurr1 activity at the same ligand concentrations tested in the Nurr1-dependent NBRE and NurRE luciferase assays (**Figure 2**) for direct comparison to assess Nurr1-dependent and Nurr1-independent effects on transcription. The first assay tests for compound toxicity and overall effects on general transcription; cells are transfected with a reporter plasmid containing five copies of the yeast Gal4 upstream activation sequence (UAS) followed by the firefly luciferase gene along with an expression plasmid containing the yeast Gal4 DNA-binding domain fused to the herpes simplex virus protein VP16 activation domain. The Gal4-VP16 fusion protein displays constitutively high luciferase activity; compounds that show decreased activity may either display cytotoxicity or inhibit general transcription in a Nurr1-independent manner, whereas compounds that show increased activity activate general transcription in a Nurr1-indepenent manner. We also tested the compounds using the CellTiter Glo luminescence cell viability assay as a more direct measure of cytotoxicity.

Amodiaquine (**Figure 3A**) and chloroquine (**Figure 3B**) activated VP16 in SK-*N*-BE(2)-C cells at lower concentrations and abruptly decreased activity at the highest concentration tested. However, the compounds only showed cytotoxicity at the highest concentration tested in HEK293T cells, indicating they affect general transcription via Nurr1-independent mechanisms. SR10098 (**Figure 3C**), SR24237 (**Figure 3D**), SR10658 (**Figure 3E**), and IP7e (**Figure 3F**) activated VP16 in the same cell lines that showed activation in the NBRE and NurRE luciferase assay without any cytotoxicity, indicating they also affect general transcription via Nurr1-independent mechanisms. C-DIM12 (**Figure 3G**) and camptothecin (**Figure 3H**), which showed decreased Nurr1 activity in some of the NBRE and NurRE assays, also showed decreased VP16 and/or CellTiter Glo activity in the same cell lines, indicating that Nurr1-independent and cytotoxic mechanisms contribute to activity of these compounds. Finally, celastrol (**Figure 3I**), isoalantolactone (**Figure 3J**), cytosporone B (**Figure 3K**), and TMPA (**Figure 3L**) showed VP16 and CellTiter Glo activity trends consistent with Nurr1-independent transcriptional mechanisms. Taken together, these data do not rule out that the ligands display Nurr1-dependent transcriptional mechanisms; however, the data indicate most if not all of the ligands also display Nurr1-independent effects on general transcription in a cell type-specific manner in the same conditions where the compounds affected the activity in the NBRE and NurRE assays.

**Fig. 3.**
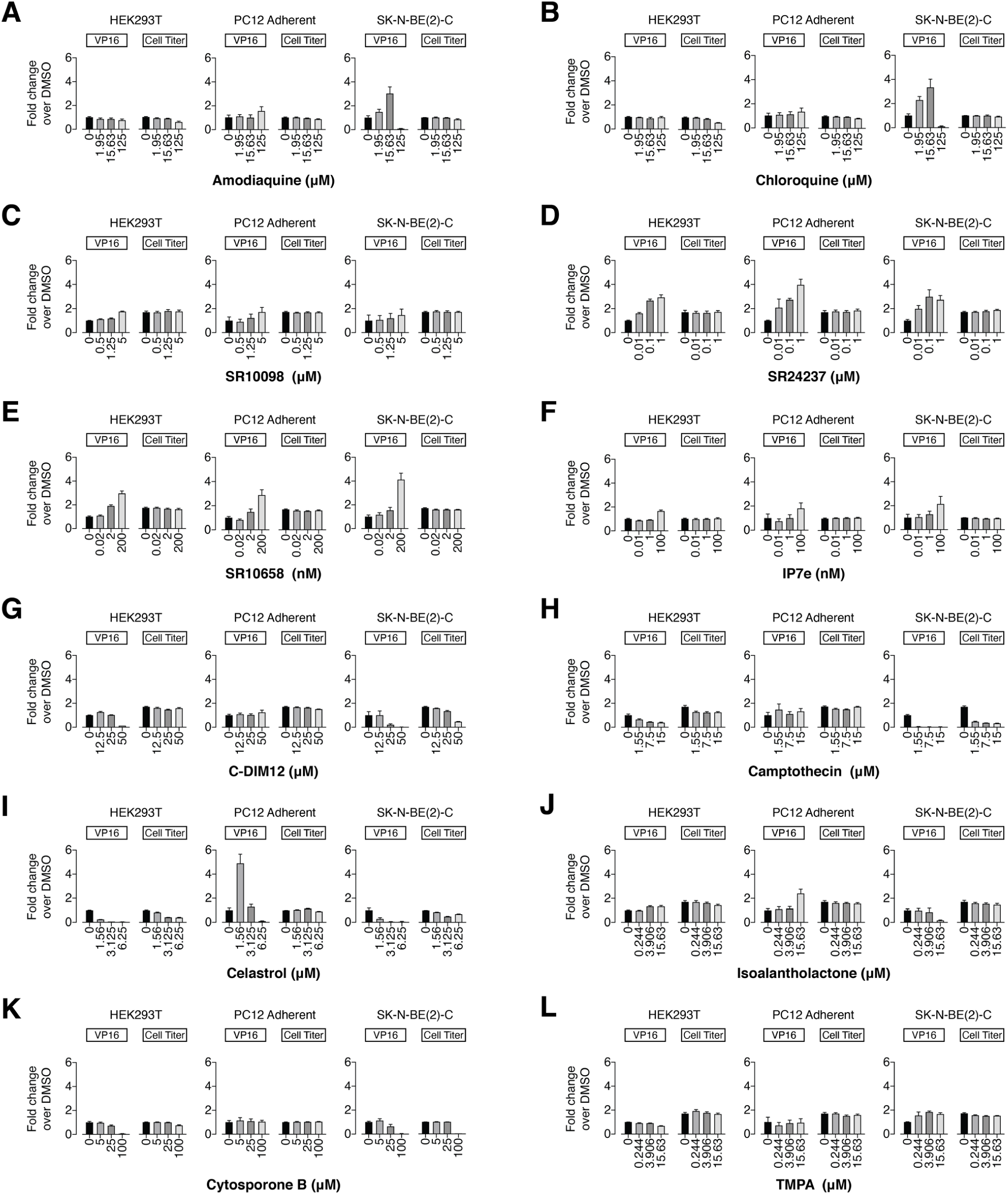
Effect of the twelve NR4A ligands on general transcription assessed using a Gal4-VP16 control fusion protein with a 5x-UAS-luciferase plasmid and cytotoxicity using a CellTiter-Glo assay in HEK293T, PC12, and SK-N-BE(2)-C cells (n=4, mean ± s.e.m; representative of ≥ 2 independent experiments).

### Protein NMR structural footprinting

The notion that the NR4A ligands show Nurr1-independent mechanism on general transcription raise a question as to whether they physically interact with and bind to the Nurr1 LBD as opposed to binding upstream effector proteins (e.g., kinases) that in turn regulate Nurr1 transcription through downstream cellular functions. We therefore used a protein NMR spectroscopy structural footprinting assay to determine if the NR4A ligands directly bind to the Nurr1 LBD. We collected 2D [^1^H,^15^N]-TROSY-HSQC of ^15^N-labeled Nurr1 LBD in the presence of vehicle control or 2 molar equivalents of ligand (Figure 4). We then calculated the NMR chemical shift perturbation (CSP) (Figure 5A) and NMR line broadening via peak intensity changes (Figure 5B). Consistent with previous data (*4, 11*), addition of amodiaquine and to a lesser degree chloroquine, which is less potent than amodiaquine (*11*), showed select NMR CSP and peak intensity changes for Nurr1 LBD residues within helix 3, helix 6, helix 10/11 and helix 12, indicating they directly bind to the Nurr1 LBD. Of the other ten NR4A ligands, only cytosporone B showed NMR structural footprinting results indicating direct binding to the Nurr1 LBD; SR10098, SR24237, SR10658, IP7e, C-DIM12, camptothecin, celastrol, isoalantolactone, and TMPA showed no evidence of binding. The profile of cytosporone B-induced CSP and peak intensity changes are similar to the changes caused by amodiaquine, indicating they share a common binding site within the Nurr1 LBD.

**Fig. 4.**
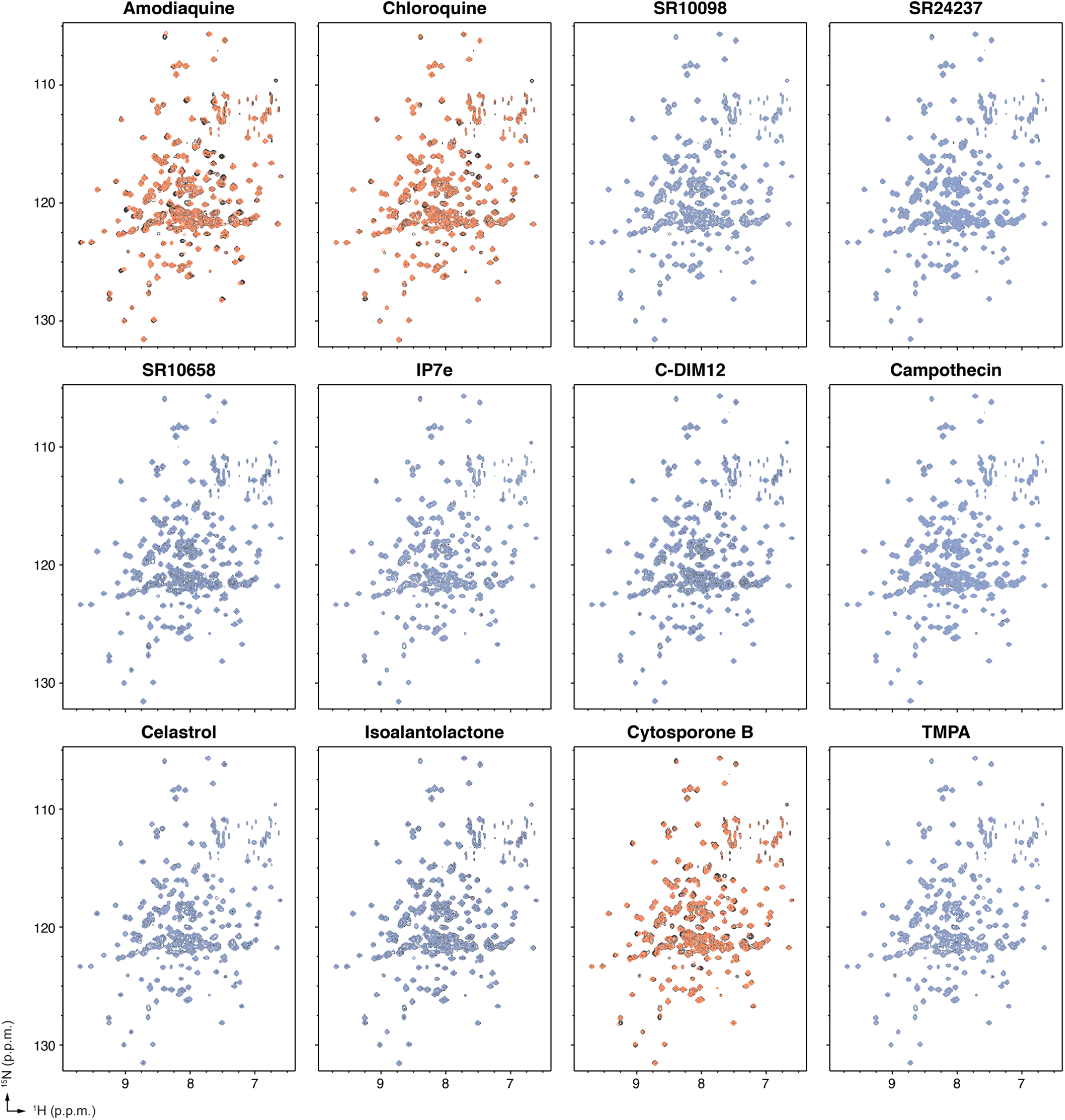
Protein NMR spectroscopy ligand footprinting data. 2D [^1^H,^15^N]-TROSY-HSQC data of ^15^N-labeled Nurr1 LBD in the absence (black spectra) or presence of 2X ligand reveals ligands that do not bind to the Nurr1 LBD (blue spectra) and ligands that bind to the Nurr1 LBD (orange spectra).

**Fig. 5.**
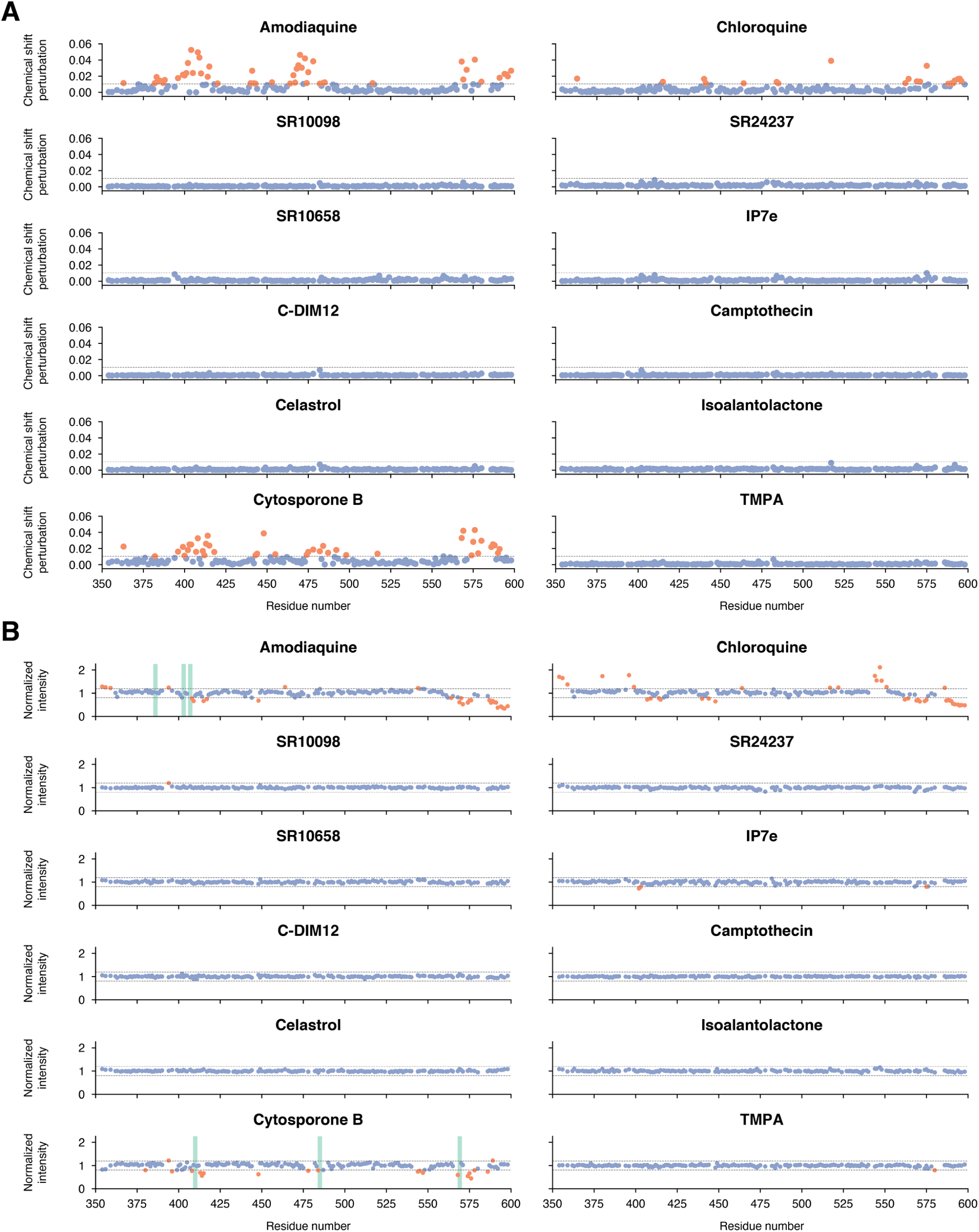
Quantitative analysis of the 2D NMR ligand footprinting data, including (**A**) differential NMR chemical shift perturbation analysis of DMSO vs. 2X ligand and (**B**) NMR peak intensity analysis showing residues with no significant effect (blue circles), residues with a CSP or peak intensity change ≥ mean ± 4 s.d. (indicated by a dotted line; orange circles), or residues with peak intensities that broadened or decreased into the noise (green bars).

To visualize the NMR-detected binding effects, we mapped the NMR structural footprinting changes for amodiaquine, chloroquine, and cytosporone B, onto the crystal structure of Nurr1 LBD (*2*) and compared the binding epitopes to the crystal structure of TMPA-bound Nur77 LBD (*30*), PDNPA-bound Nur77 LBD (*31*), and the Nur77 modeled binding sites of cytosporone B (*29*) and celastrol (*25*) relative to the canonical ligand-binding pocket (Figure 6). Amodiaquine, chloroquine, and cytosporone B show similar binding epitopes on the Nurr1 LBD, which most of the NMR-detected changes occurring for residues within or near the canonical orthosteric-ligand binding pocket. For Nur77, the crystallized TMPA binding poses and modeled celastrol and cytosporone B binding poses are all surface exposed.

**Fig. 6.**
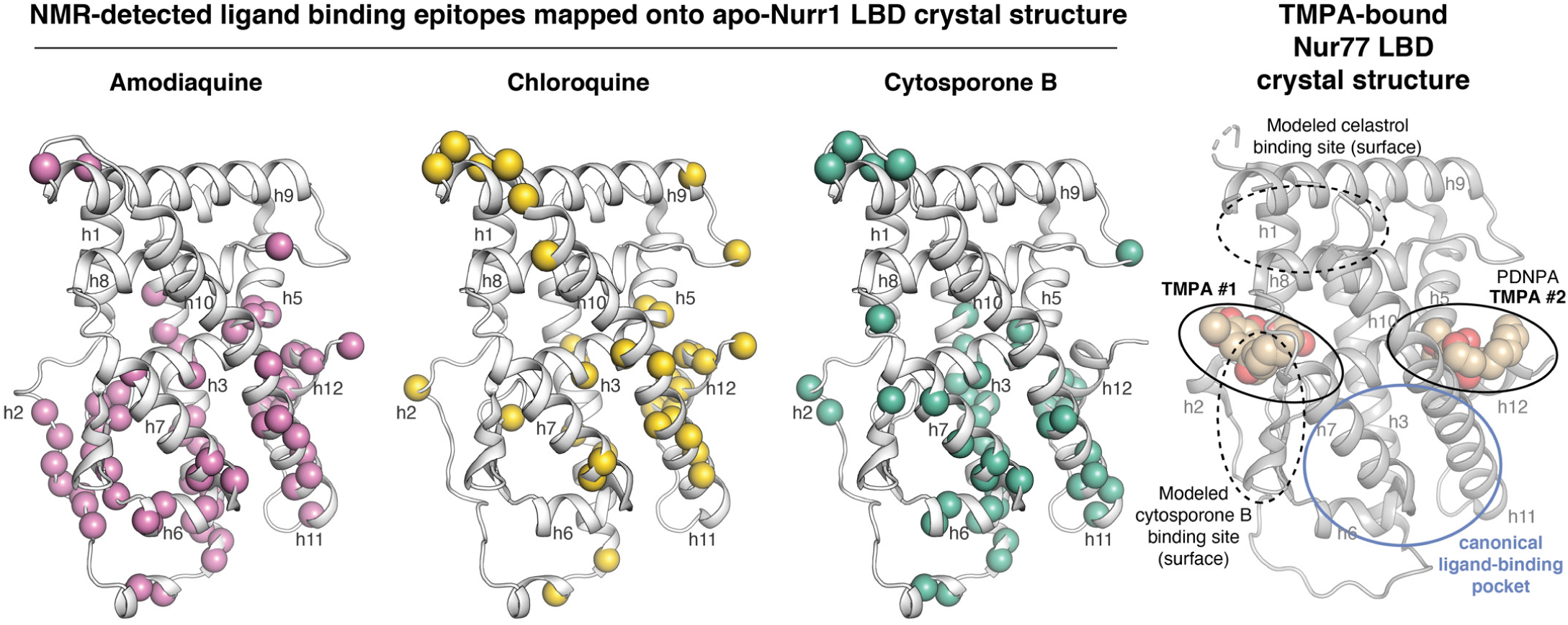
Mapping the NMR structural footprinting data of the binding ligands onto the Nurr1 LBD crystal structure (PDB 1OVL, chain B) and comparison to TMPA-bound Nur77 LBD (PDB 3V3Q) along with other modeled and crystallized Nur77 ligand binding sites reported in the literature. Spheres indicate residues with NMR CSP or peak intensity values with large changes (mean ± 4 s.d.) in the 2D NMR ligand footprinting data.

Although there is some overlap between the surface exposed Nur77 interaction sites and the Nurr1 NMR-detected binding epitopes, there are also differences. For example, cytosporone B and TMPA are derived from the same scaffold, and the modeled cytosporone B interaction site agrees with one of the two crystallized TMPA binding modes—a solvent exposed surface in the Nur77 LBD. However, the NMR-detected cytosporone B binding epitope on Nurr1 is different, suggesting the interaction occurs within the canonical ligand-binding pocket similar to amodiaquine, chloroquine, and unsaturated fatty acids (*4, 5, 11*). These data suggest that cytosporone B likely binds differently to Nurr1 and Nur77, but another explanation could be that the solution NMR structural footprinting analysis picks up on binding events or structural changes that are not apparent in solid state crystallography studies.

## DISCUSSION

Defining if and how ligands bind to Nurr1 is critical not only for understanding Nurr1 function and regulation but also in prioritizing and directing medicinal chemistry efforts on Nurr1-binding ligands. Relative to other nuclear receptors, crystal structures have not revealed a well-defined ligand-binding pocket in the Nurr1 LBD. Solution-state structural studies indicate the Nurr1 ligand-binding pocket is dynamic and solvent accessible, indicating the absence of a pocket captured in Nurr1 LBD crystal structures represents a collapsed conformation (*4*). The lack of a well-defined Nurr1 ligand binding pocket has arguably stunted efforts to discover and design Nurr1-binding compounds. Several compounds have been reported in the literature to interact with the Nurr1 LBD. Unsaturated fatty acids, amodiaquine, and chloroquine appear to bind to the Nurr1 orthosteric ligand-binding pocket (*4, 5, 11*), whereas the endogenous dopamine metabolite 5,6-dihydroxyindole (DHI) covalently binds to noncanonical site via covalent attachment to a surface-exposed cysteine residue in the Nurr1 LBD (*7*).

Our studies here show that most of the other NR4A ligands that influence the cellular functions of Nurr1 do not appear to function through direct binding to Nurr1. These findings indicate that these ligands exert their effects on Nurr1 activity through binding to upstream effector proteins such as kinases, which could then affect Nurr1 cellular activity via downstream effects. Several other observations support this idea.

First, most if not all of the NR4A ligands show cell type-specific functions. It is possible this is due to the availability of different transcriptional coregulator proteins present within different cell types that could be recruited to Nurr1 in a ligand-dependent manner. However, given that most of the NR4A ligands do not directly bind Nurr1, it is possible that cell type-specific expression of upstream effector proteins also contributes to the cell type-specific activities of the ligands. Cell type-specific Nurr1 activity has been highlighted previously in neuronal cell lines, suggesting that endogenous factors expressed in specific neuronal cell lines influence Nurr1 activation (*32*). Cell type-specific dependence may apply to other types including bladder cancer cells and other human cancer cells where Nurr1 activity has been found to be critical for survival (*33-35*).

Second, some of these NR4A ligands are polypharmacology modulators. As one example, C-DIM12 and related analogs were reported as Nurr1 activators in pancreatic cancer cells (*21*), bladder cancer cells (*34*), and neuronal cells (*22, 23*); a Nurr1 inhibitor in glioblastoma cells (*36*); and activates other nuclear receptors in various cell types including Nur77 (*37, 38*), COUP-TF1 (*39*) and PPARγ (*40-42*). Indeed, previous studies suggest that the mechanisms of action of these C-DIM compounds may occur independent of nuclear receptor binding or via nuclear receptor-independent mechanisms through affectingkinase activity (*38, 43-48*). *In silico* ligand docking studies suggested that C-DIM12 may bind to the Nurr1 LBD coregulator binding surface (*23*). However, our protein NMR structural footprinting data clearly show C-DIM12 does not directly bind to the Nurr1 LBD. Further selectivity profiling such as chemoproteomic methods is warranted to determine the molecular target of C-DIM12 and related analogs.

Related to polypharmacology, some of the other NR4A ligands that we profiled that do not directly bind to the Nurr1 LBD but activate Nurr1 transcription have been shown to function through targets other than the NR4As, which in principle could affect Nurr1 activity through down-stream functions of the targets. Celastrol, which contains a reactive quinone methide moiety enabling covalent attachment to cysteine residues (*49*), including direct binding and inhibition of c-Myc-Max heterodimers (*50*), cancerous inhibitor of protein phosphatase 2A (CIP2A) (*51*), STAT3 (*52*), SHOC-2 to inhibit ERK signaling (*53*), IKKα and IKKβ (*54*), and HSP90-chaperone interactions (*55-57*); directly binds nearly 70 protein targets in a proteome microarray assay (*53*); and affects other cellular signaling pathways including protein phosphatase 2A-Akt, AMPK, and WNT/β-catenin (*58*). Isoalantolactone activates AMPKα (*28*) and inhibits STAT3 (*59*) and IKKβ (*60*). Camptothecin inhibits topoisomerase I (*61*). Given that the other Nurr1 activators identified in HTS screens or derived from HTS hits (SR10098, SR10658 and the related analog IP7e, and SR24237) do not directly bind to the Nurr1 LBD, these compounds likely target other effector proteins that influence Nurr1 activity or other general transcriptional machinery since some showed activity in the VP16-Gal4 assay. This concept of ligands affecting Nurr1 activity via binding to upstream effectors of Nurr1 is supported by a study showing that kinase inhibitors can activate and inhibit Nurr1 transcription (*62*). Thus, kinases could act as upstream effector proteins on downstream Nurr1 activities.

Amodiaquine and chloroquine were two of the three compounds in our panel of twelve NR4A ligands that physically bind to the Nurr1 LBD and activated Nurr1 transcription in a screen using SK-*N*-BE(2)-C neuronal cells (*11*). However, we also found that amodiaquine and chloroquine activated transcription in the VP16-Gal4 assay in SK-*N*-BE(2)-C cells, indicating they also have Nurr1-independent effects on transcription. Amodiaquine is well known as an antimalarial drug mostly used against strains of *Plasmodium falciparum* (*63*), but it also inhibits human histamine N-methyltransferase (*64*) and several human cytochrome P450 enzymes (*65*). Chloroquine, which is also an antimalarial drug, is a chemokine receptor CXCR4 antagonist (*66*) and used as anticancer agent capable of inhibiting autophagy by disrupting the fusion of autophagosomes with lysosomes (*67, 68*). Future work on this 4-aminoquinoline scaffold may result in the development of direct Nurr1-binding compounds with better specificity towards Nurr1 and reduced general effects on transcription.

Our NMR studies show that cytosporone B, which was the first identified Nur77 agonist (*29*)—but not the related analog TMPA (*30*)— directly binds to the Nurr1 LBD. We found that cytosporone B did not activate Nurr1 transcription in HEK293T cells or the PC12 and SK-*N*-BE(2)-C neuronal cell lines. However, it is possible that cytosporone B displays cell type-specific activities, as cytosporone B was previously shown to activate Nur77 and Nurr1 in BGC-823 human gastric cancer cells (*29*). Future studies are needed to detail whether cytosporone B can affect Nurr1 transcription through direct binding in other cell types where Nurr1 is expressed to determine if it represents a potential starting point for future medicinal chemistry efforts on Nurr1-binding ligands.

In conclusion, our studies emphasize the need for determining whether ligands that affect Nurr1 activity, or NR4A activity more broadly, indeed bind directly to Nurr1 or function through upstream effector proteins. Future studies employing biochemical, biophysical, or structure-based screening might lead to new direct binding Nurr1/NR4A ligand scaffolds, which would open new possibilities for developing Nurr1-binding ligands.

## MATERIALS AND METHODS

### Compounds and chemical synthesis

Nine of the twelve NR4A compounds were purchased from commercial vendors: amodiaquine (Xenotech Llc), chloroquine (Chem Impex Intl Inc), IP7e (Tocris), C-DIM12 (Sigma-Aldrich), camptothecin (Cayman), celastrol (Cayman), isoalantolactone (Indofine), cytosporone B (Tocris), and TMPA (EMD Millipore). Three compounds were synthesized in-house using previously described methods including SR10098 (*17*), SR10658 (*18*), and SR24237 (*20*); purity (>95%) was confirmed using an Agilent 1100 series HPLC system and identity was confirmed by ^1^H NMR using a Bruker 600 MHz NMR spectrometer and mass analysis using a Thermo Scientific Ultimate 3000/LCQ Fleet system (ESI) mass spectrometer. Compounds were suspended according to vendor recommendations when applicable in water (chloroquine), ethanol-d_6_ (amodiaquine), or DMSO-d_6_ (all other ligands).

### Spectral characterization of synthesized compounds

**1 (SR10098)**. ^1^H NMR (DMSO-D_6_, 600 MHz), δ(p.p.m.): 8.42-8.4 (m, 1H), 8.23-8.19 (m, 2H), 8.16 (dd, 1H), 7.79 (dd, 1H), 7.48-7.43 (m, 2H). MS (ESI): Expected mass for C14H9FN2O2 (M + H)^+^: 256.06 Da, observed mass: 256.79 Da.

**2 (SR24237)**. ^1^H NMR (DMSO-D_6_, 600 MHz), δ(p.p.m.): 10.37 (s, 1H), 9.49 (s, 1H), 8.79-8.77 (m, 1H), 8.69 (d, 1H), 8.18 (dd, 1H), 8.09 (dt, 1H), 8.03 (dt, 1H), 7.98-7.96 (m, 2H), 7.83 (dt, 1H), 7.49 (ddd, 1H), 7.43-7.39 (m, 1H), 7.18-7.15 (m, 1H). MS (ESI): Expected mass for C19H14N4O (M + H)^+^: 314.12 Da, observed mass: 314.80 Da.

**3 (SR10658)**. ^1^H NMR (DMSO-D_6_, 600 MHz), δ(p.p.m.): 8.17-8.15 (m, 2H), 7.53-7.44 (m, 5H), 7.06-7.03 (m, 2H), 6.74 (s, 1H). MS (ESI): Expected mass for C26H22N2O3 (M + H)^+^: 334.13 Da, observed mass: 332.87 Da.

### Cell lines

All cell lines were obtained from ATCC, including HEK293T (#CRL-11268), PC12 (#CRL-1721.1), and SK-*N*-BE(2)-C (#CRL-2268) and cultured according to ATCC guidelines. HEK293T cells were grown at 37 °C, 5% CO_2_ in DMEM (Gibco) supplemented with 10% fetal bovine serum (Gibco) and 100 units/mL of Penicillin, 100 µg/mL of Streptomycin and 0.292 mg/mL of Glutamine (Gibco) until 90 to 95% confluence prior to subculture or use. SK-*N*-BE(2)-C were maintained at 37 °C, 5% CO_2_ in a media containing 1:1 mixture of EMEM (ATCC) and F12 medium (Gibco) supplemented with 10% fetal bovine serum (Gibco) until 90 to 95% confluence prior to subculture or use. PC12 cells were grown at 37 °C, 5% CO_2_ in FK-12 medium (Gibco) supplemented with 15% horse serum (Sigma) and 2.5% fetal bovine serum (Gibco) until 90 to 95% confluence prior to subculture or use.

### Cellular transcription assays

HEK293T (#CRL-11268), PC12 (#CRL-1721.1), and SK-*N*-BE(2)-C (#CRL-2268) cells were seeded in 10-cm petri dish at 1.5 million cells. The following day, cells were transfected using Lipofectamine 2000 (Thermo Fisher Scientific) and Opti-MEM with full-length Nurr1 expression plasmid (2 µg), 3xNBRE3 or 3xNurRE-luciferase reporter plasmid (6 µg), to a total of 8 µg total DNA and incubated for 18 h. For Gal4-VP16 transactivation, cells were transfected the same way but with a Gal4-VP16 expression plasmid (2 µg) and 5xUAS-luciferase reporter plasmid (Upstream Activation Sequence; 2µg). Cells were transferred to a white 384-well plates (Thermo Fisher Scientific) at 10,000 cells/well in 20 µL and incubated for 4 h. Ligands were prepared in dose response dilutions using vehicle control: water (chloroquine), ethanol (amodiaquine), or DMSO (all others). Ligand in dose response format or the respective vehicle control were added to the cells (20 µL). Cells were then incubated for 18 h and harvested for luciferase activity quantified using Britelite Plus (Perkin Elmer; 20 µL) on a Synergy Neo plate reader (Biotek). Data were plotted as bars and analysis performed using GraphPad Prism.

### Cell viability assays

HEK293T, PC12 or SK-N-BE(2)-C cells were seeded in 10-cm petri dish at 1.5M cells. The following day, Cells were transferred to a white 384-well plates (Thermo Fisher Scientific) at 10,000 cells/well in 20 µL and incubated for 4 h. Ligands (or vehicle control) were added (20 µL), cells incubated for 18 h and harvested for cell viability quantitation using Cell-Titer-Glo (Promega; 20 µL) on a Synergy Neo plate reader (Biotek).

### Expression and purification of ^15^N-labeled Nurr1 LBD

Recombinant ^15^N-labeled Nurr1 LBD (NR4A2; residues 353 to 598) was expressed and purified as previously described (*5*). Briefly, the protein was expressed in *Escherichia coli* BL21(DE3) cells (Life Technologies) using a pET-46 tobacco etch virus (TEV) protease-cleavable N-terminal hexahistidine tag fusion protein in M9 media supplemented with ^15^NH_4_Cl (Cambridge Isotope Labs, Inc.). Nurr1 LBD was eluted against a 500 mM imidazole gradient through a Ni-NTA column, followed by overnight dialysis against a buffer without imidazole for TEV protease His tag cleavage at 4 °C. The next morning, the sample is loaded onto the Ni-NTA column for contaminants and tag removal. The flow through containing the purified protein was collected, concentrated and ran through a S75 size exclusion column (GE healthcare) in NMR buffer (20 mM KPO_4_ pH 7.4, 50 mM KCl, and 0.5 mM EDTA). The corresponding protein peak is collected and stored at −80 °C. All the ligands were dissolved in either water, DMSO-d_6_, or ethanol-d_6_ for NMR experiments.

### Protein NMR spectroscopy structural footprinting

Data were collected on a Bruker 700 MHz NMR spectrometer equipped with a QCI cryoprobe at 298 K. For each ligand titration, 2D [^1^H,^15^N]-TROSY-HSQC were acquired at 298°K using 200 µM ^15^N-labeled Nurr1 LBD in the absence or presence of 400 µM ligand in NMR buffer containing 10% D_2_O. Data were processed using NMRFx (*69*) and analyzed using NMRViewJ (*70*). Chemical shift perturbation (CSP) analysis was performed by transfer of Nurr1 LBD NMR chemical shift assignments (*71*) that we previously validated and reported (*5*) from the vehicle to the 2X ligand spectra using the minimal NMR chemical shift method (*72*). Peaks were identified to have broadened to zero if there was no confident peak in proximity to the vehicle peak. The average CSP and the standard deviation (SD) in the CSPs was calculated for titrations after rejecting outliers more than 2 SD from the mean, and the peaks that displayed CSPs in the presence of peptide greater than ≥4 SD above the average CSP were noted as significant.

## ACKNOWLEDGEMENTS

This work was supported in part by National Institutes of Health (NIH) grants R01GM114420 (D.J.K.) and S10OD021550 (which along with institutional funds from Scripps Research funded purchase of the Bruker 600 MHz NMR).

## AUTHOR CONTRIBUTIONS

P.M.T. and D.J.K conceived and designed the research. P.M.T. performed cellular assays and protein NMR analyses. H.L, P.K, and T.M.K. synthetized compounds. I.M.S.dV. contributed to preliminary NMR studies on this project. D.J.K. analyzed data and supervised the research. P.M.T. and D.K. wrote the manuscript with input from all authors. The authors collectively declare no conflicts of interest in the completion of this study.

